# Parallel mechanisms detect different photoperiods to independently control seasonal flowering and growth in plants

**DOI:** 10.1101/2023.02.10.528016

**Authors:** Qingqing Wang, Wei Liu, Chun Chung Leung, Daniel A. Tartè, Joshua M. Gendron

## Abstract

For nearly 100 years, we have known that both growth and flowering in plants are seasonally regulated by the length of the day (photoperiod). Intense research focus and powerful genetic tools have propelled studies of photoperiodic flowering, but far less is known about photoperiodic growth, in part because tools were lacking. Here, using a new genetic tool that visually reports on photoperiodic growth, we identified a seasonal growth regulation pathway, from photoperiod detection to gene expression. Surprisingly, this pathway functions in long days but is distinct from the canonical long day photoperiod flowering mechanism. This is possible because the two mechanisms detect the photoperiod in different ways: flowering relies on measuring photoperiod by directly detecting duration of light intensity while the identified growth pathway relies on measuring photosynthetic period indirectly by detecting the duration of photosynthetic metabolite production. In turn, the two pathways then control expression of genes required for flowering or growth independently. Finally, our tools allow us to show that these two types of photoperiods, and their measurement systems, are dissociable. Our results constitute a new view of seasonal timekeeping in plants by showing that two parallel mechanisms measure different photoperiods to control plant growth and flowering, allowing these processes to be coordinated independently across seasons.

## Main text

Photoperiod (daylength) is a stable indicator of the season, and many organisms have evolved photoperiod measuring systems to predict abiotic and biotic changes associated with a given season ^1^. Plants have served as a preeminent study system for photoperiodism because of their propensity to flower in specific seasons, and a pathway for seasonal flowering is known. Red and blue light are sensed by photoreceptors that control the stability of the CONSTANS (CO) transcription factor which in turn activates expression of the florigen gene, *FLOWERING LOCUS T* (*FT*), which triggers the transition to flowering ^2–5^. Despite intense focus on flowering, research from more than 100 years ago shows that both flowering and growth are under the control of photoperiod in plants but are separable and can be timed differentially throughout the year. For instance, plants often grow rapidly in long days, but can flower faster in short days ^6^. However, little is known about the molecular determinants that support photoperiodic growth, including the photoperiod measuring systems, cellular signaling pathways, and genes required for photoperiod-dependent growth. This deficiency is partially due to a lack of genetic tools and markers for growth that are akin to the tools available for the CO/FT pathway.

To overcome this deficiency, we analyzed published RNA-seq and microarray data for genes that are required for long day-specific rapid growth in Arabidopsis and that also have higher expression in long days ^7–10^. One of the identified genes, *MIPS1* is required for production of myoinositol which serves as the chemical precursor for a wide variety of signaling molecules, vitamins, and membrane components required for cell division, cell expansion, and maintenance of cell viability, all important facets of vegetative growth ^11–14^. Additionally, *MIPS1can* participate in pathogen defense, another cellular process that is known to alter growth ^15,16^. We first plotted the expression of *MIPS1* from microarray and RNA-seq experiments and tested the observed expression pattern using qRT-PCR (Fig. 1a-c). *MIPS1* expression is qualitatively and quantitatively different in 16 hours light:8 hours dark (16L:8D) and 8L:16D. In 8L:16D *MIPS1* expression is monophasic, with a peak in expression at ZT4 (4 hours after dawn). In 16L:8D, *MIPS1* expression is biphasic, containing the ZT4 peak seen in 8L:16D and a second photoperiod-specific peak phased to dusk at ZT16. This results in a rDEI_16L:8D/8L:16D_ (relative Daily Expression Integral ^8,9^) of 1.44 in the microarray experiment, 2.77 in the RNA-seq experiment, and 1.70 in the qRT-PCR. These results indicate that plants grown in 16L:8D have higher *MIPS1* expression due to a photoperiod-specific peak in expression at ZT16.

**Fig. 1:**
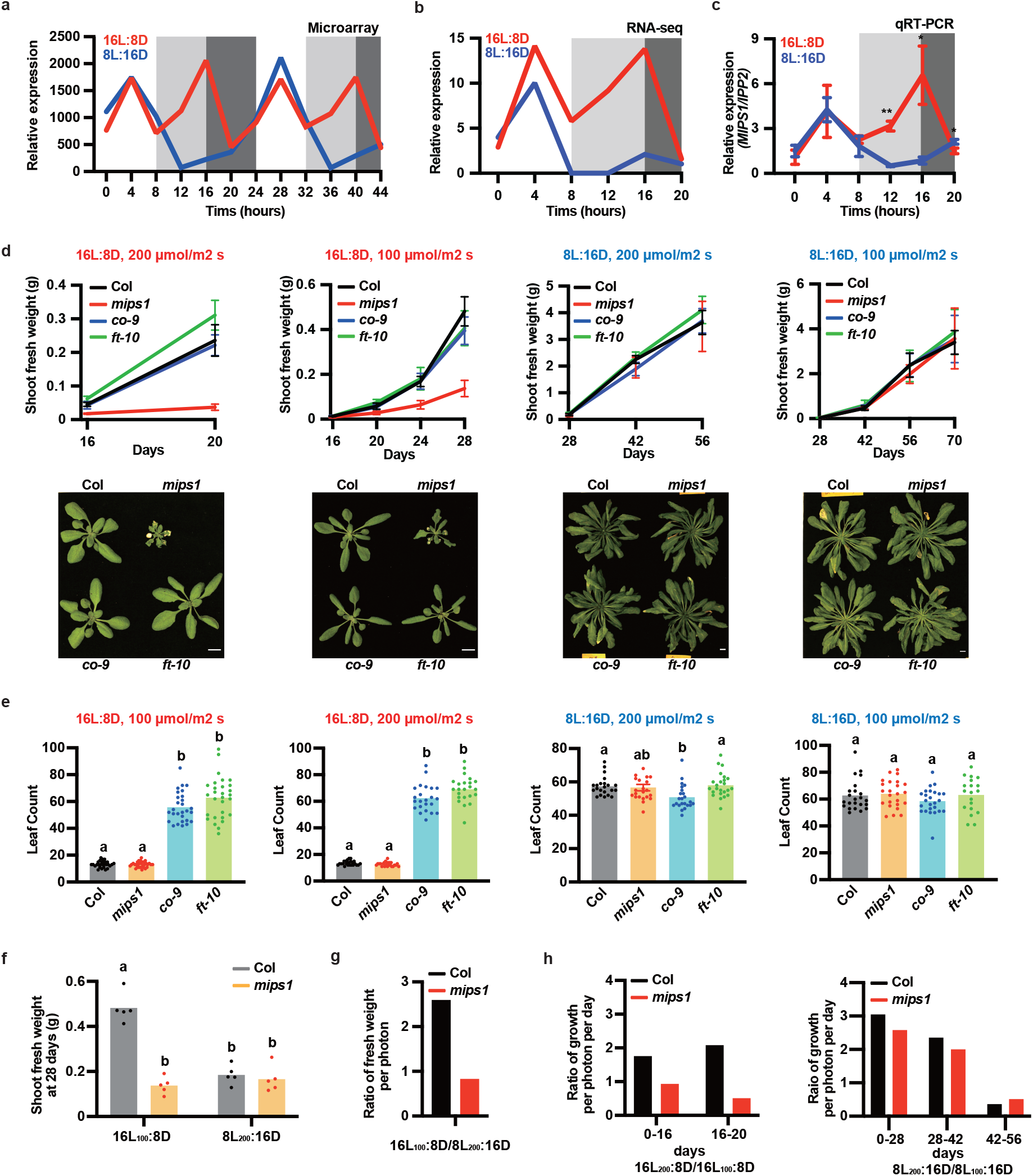
*MIPS1* is involved in photoperiod-dependent growth. **a-c,** Microarray (a), RNA-seq (b) and qRT-PCR (c) of *MIPS1* expression from 12-day-old plants grown in 16L:8D (red) and 8L:16D (blue). Shading indicates dark time. *IPP2* was used as internal control for panel c. Error bar indicates SD. *, *p* ≤ 0.05; **, *p* ≤ 0.01 (Welch’s t-test). n = 3. **d-e,** Shoot fresh weight with representative images (d) and leaf number at 1 cm bolting (e) of Col, *mips1, co-9*, and *ft-10* mutants grown in 16L:8D or 8L:16D with light intensity of 100 μmol/m^2^ s or 200 μmol/m^2^ s as indicated. Error bars indicate SD (n = 5 in d and n = 19 - 30 in e). Scale bar = 1 cm. **f,** Shoot fresh weight of Col and *mips1* from 28-day-old plants grown in 16L:8D or 8L:16D condition with light intensity of 100 μmol/m^2^ s or 200 μmol/m^2^ s as indicated. n = 5. **g,** Ratio of growth per available photon of 28-day-old Col and *mips1* plants grown in16L_100_:8D compared to 8L_200_:16D. Growth per available photon = Shoot fresh weight / days of growth / photon per day. **h,** Ratio of growth per available photon of Col and *mips1* plants grown in16L_200_:8D compared to16L_100_:8D and 8L_200_:16D compared to 8L_100_:16D at indicated age. Growth per photon = shoot fresh weight / days of growth / photon per day. In e and f, different letters indicate significant differences as determined by one-way ANOVA followed by Dunnett’s T3 multiple comparison test; p≤0.05. Raw data and calculations for physiology experiments are presented in Supplementary Table 2.

Previously, *mips1* mutant plants were reported to have growth defects that are dependent on the daily light integral (the amount of photosynthetically active light passing through a defined area in 24 hours) and possibly photoperiod, although these defects varied based on experimental conditions ^11,17^. We wanted to distinguish the role, if any, that *MIPS1* has on photoperiodic growth. Thus, we designed a multifactorial growth experiment that allowed us to distinguish between intensity-specific and photoperiod-specific effects on growth. We germinated and grew wild-type and *mips1-2* (SALK_023626) mutant plants in two light intensities (100 and 200 μmol/m^2^ s) and two photoperiods (16L:8D and 8L:16D) and measured pre-flowering vegetative fresh rosette weight as a proxy for growth. We collected samples over the vegetative growth phase of the plant (Fig. 1d).

In this experiment, the *mips1-2* mutant is unable to generate fresh weight at the same rate as wild type in the 16L:8D growth conditions but has no observable fresh weight defect in the 8L:16D growth conditions. To determine the role of *MIPS1* in photoperiod-dependent growth, independent of the daily light integral, we compared the plants grown in 16L:8D in 100 μmol/m^2^ s white light (16L_100_:8D) and 8L_200_:16D on day 28 (Fig. 1f, g and Supplementary Fig.1e, g). In these conditions the daily light integral is the same, but the wild-type plants grown in 16L_100_:8D have generated 2.6 times the mass of the 8L_200_:16D-grown plants, confirming the role of photoperiod in Arabidopsis growth.

By dividing the fresh weight by the daily light integral and then calculating a ratio between two conditions, we can calculate relative fresh weight gain per available photon. This is a simple measure of how well each plant can utilize available light for growth, but it must be considered that this value is affected by leaf shape, leaf thickness, and other physiological factors. Despite these constraints, this value can help understand how well the plants are using available light in different photoperiods and light intensities for generation of fresh weight. For the comparison of the plants in 16L_100_:8D and 8L_200_:16D, the daily light integral is the same between the two conditions, thus the result is the same value as the fold change in mass and shows that in 16L_100_:8D the plants are using each available photon 2.6 times more effectively for growth than in 8L_200_:16D (Fig. 1g). Conversely, the *mips1-2* mutant plants grown in 16L_100_:8D have generated 0.83 times the mass of the 8L_200_:16D-grown plants, demonstrating that the *mips1-2* mutant has lost the ability to generate rosette fresh weight in a photoperiodic manner.

To determine the role of *MIPS1* in intensity-dependent growth, we can compare rosette fresh weight when light intensity is doubled in 16L:8D or 8L:16D (Fig. 1h and Supplementary Fig.1f). Wild-type plants in 16L_200_:8D are 3.5 or 4 times larger than in 16L_100_:8D at day 16 and 20, respectively, meaning they can generate up to twice as much fresh weight per available photon, and this only slightly changes between plants that are 16 or 20 days old. Plants in 8L_200_:16D can be 6.1, 4.8 or 1.5 times larger than plants grown in 8L_100_:16D, which shows that early in development they can generate up to three times as much weight per additional photon, but this effective use of the increased photons for growth decreases as the plants age. In 16L:8D the *mips1-2* mutant has lost the ability to generate rosette fresh weight in an intensity-dependent manner, but this ability is unaffected in 8L:16D. This indicates that wild-type plants alter intensity-dependent growth responses based on the photoperiod and that *MIPS1* plays a role in these photoperiod-specific growth processes.

### *MIPS1* is required for leaf growth and reproductive fitness, independent of flowering time

In addition to fresh weight, we also measured flowering time using two metrics, number of days to reach a 1 cm floral bolt and number of leaves at the time when the floral bolt reaches 1 cm (Fig. 1e and Supplementary Fig.1a). Despite severe growth defects in 16L:8D, the *mips1-2* mutant has no measurable defect in flowering time in 16L:8D or 8L:16D. Identical leaf count numbers between wild type and *mips1-2* mutants, in combination with the rosette fresh weight data, indicate that *MIPS1* has no role in flowering or leaf initiation rate during the normal vegetative growth period but is required for generating leaf mass in 16L:8D.

Post-flowering development was also assessed by elongation rate of the inflorescence stem (days from one to ten cm inflorescence stem length), anthesis (first observable flower opening), and seed yield (total mass of dry seeds) (Supplementary Fig.1b, c). In wild-type plants, elongation rate of the stem is lower in 16L:8D than 8L:16D, but it is not affected by intensity in either photoperiod, while anthesis is faster in 16L:8D than 8L:16D. The *mips1-2* mutant delays both developmental processes in 16L:8D but not 8L:16D, with the exception of anthesis in 16L_100_:8D where *mips1-2* is not different than the wild type.

We used seed yield as a measure of the fitness and found that wild-type yield is greater in 16L:8D than 8L:16D and that doubling light intensity can increase yield in 16L:8D but not 8L:16D (Supplementary Fig.1d). In 16L:8D the *mips1-2* mutant has lower yield compared to the wild type in both light intensities. Conversely, we detected no yield difference between the wild type and *mips1-2* in 8L:16D at both light intensities. Thus, we conclude that *MIPS1* is required for long day-specific pre-flowering vegetative growth, post-flowering bolt elongation, and reproductive fitness but has no effect on flowering time or leaf organogenesis. The easily observable photoperiod-dependent growth defect of the *mips1* mutant makes it ideal for studying how plants measure photoperiod to support rapid long day growth.

### *MIPS1* photoperiodic expression is independent of CO

The photoperiod-controlled expression peak of *MIPS1* is phased to the same time of day (ZT16) as the expression peak for *FT* in 16L:8D ^2^, making CO a candidate regulator of *MIPS1* and photoperiodic growth. We assayed photoperiodic growth in the *co-9* ^18^ and *ft-10* ^19^ mutants which have compromised photoperiodic flowering and saw no defects in fresh weight (Fig. 1d, e). Interestingly, the *co-9* and *ft-10* mutants had reduced seed yield in 16L:8D and *ft-10* had reduced yield in 8L_100_:16D (Supplementary Fig. 1d). We then tested whether *MIPS1* expression is altered in the *co-9* mutant. We performed qRT-PCR with samples from wild-type and *co-9* mutant plants collected at ZT12 in 16L:8D and 12L:12D, a time at which *FT* is normally induced. We found *FT* expression is reduced in the *co-9* mutant, while *MIPS1* expression is not altered (Fig. 2a). Next, we generated a *co-9 mips1-2* double mutant. The double mutant shows no additional growth defects beyond those of the *mips1-2* mutant alone in 16L_100_:8D condition (Fig. 2b). Interestingly, the *co-9 mips1-2* double mutant requires more days to bolt than the *co-9* mutant, although leaf numbers are the same at time of bolting in the 16L_100_:8D condition (Fig. 2b and Supplementary Fig. 2a). This suggests that *MIPS1* may play a small role in promotion of leaf development in the absence of *CO*.

**Fig. 2:**
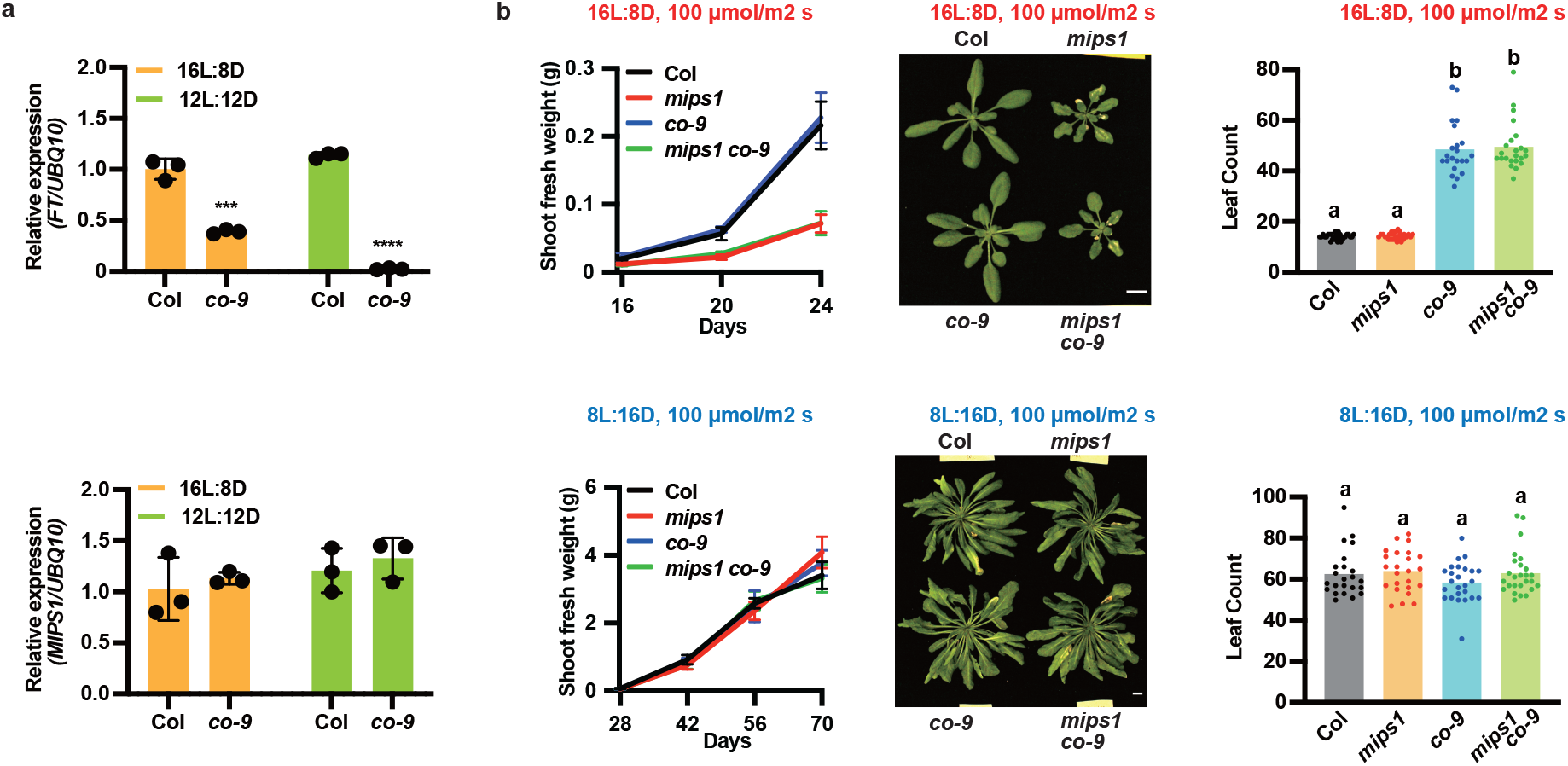
*MIPS1* expression and function are distinct from CO/FT. **a,** qRT-PCR of *FT* and *MIPS1* from 12-day-old wild-type (Col) and *co-9* mutant grown in 16L:8D (red) and 12L:12D (green) condition. Samples were harvested at ZT12. n = 3. Error bar indicates SD. ***, *p*≤0.001; ****, *p*≤0.0001 (Welch’s t-test). **b,** Shoot fresh weight with representative images and leaf number at 1 cm bolting of Col, *mips1*, *co-9*, and *mips1 co-9* mutants grown in 16L:8D or 8L:16D condition with light intensity of 100 μmol/m^2^ s. Different letters indicate significant differences as determined by one-way ANOVA followed by Dunnett’s T3 multiple comparison test; p≤0.05. Error bars indicate SD (n = 3 in a and n = 5-30 in b). Scale bar = 1 cm. The data for flowering time of Col, *mips1*, and *co-9* were re-plotted from Fig. 1e. Raw data is presented in Supplementary Table 2.

### The MDLM system regulates *MIPS1*

Recently, the metabolic daylength measurement (MDLM) system was shown to induce expression of genes in short-day photoperiods, but it remains to be seen if MDLM can also cause induction of gene expression in long-day photoperiods ^8,9^. This system relies on light sensing by photosynthesis and the circadian clock-controlled balance between starch and sucrose to control gene expression in a photoperiodic manner. First, we determined whether photosynthesis is required for induction of *MIPS1* in 16L:8D. Using qRT-PCR, we tested expression of *MIPS1* at ZT12 from plants grown in 16L:8D and then placed in darkness or kept in light but treated with a chemical inhibitor of photosystem II called 3-(3,4-dichlorophenyl)-1,1-dimethylurea (DCMU) (Fig. 3a). In both treatments, *MIPS1* expression decreased, suggesting that the transition from light to dark is communicated through photosynthesis to control induction of *MIPS1* in 16L:8D. To determine if the light/dark transition is communicated as a metabolic change, we treated wild-type plants with sucrose in 8L:16D and measured *MIPS1* expression at ZT12 when the plants are in the dark and *MIPS1* expression is normally low (Fig. 3b). Sucrose treatment induces expression of *MIPS1* suggesting that metabolism plays an important role in controlling *MIPS1* photoperiodic expression.

**Fig. 3:**
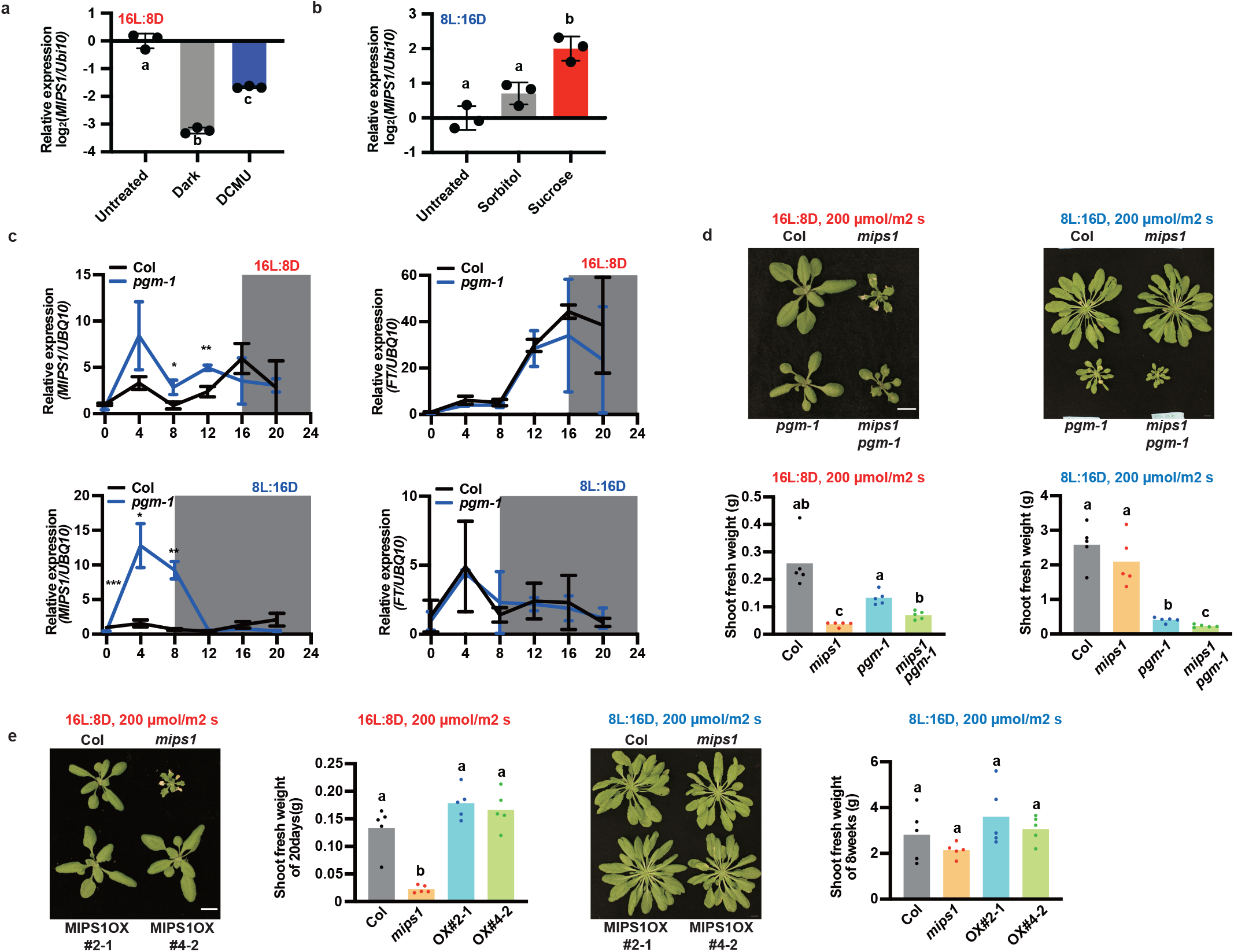
MDLM controls photoperiodic expression of *MIPS1*. **a,** qRT-PCR of *MIPS1* from 12-day-old Col plants grown in 16L:8D treated with or without darkness or DCMU at ZT0. Samples were harvested at ZT12. **b**, qRT-PCR of *MIPS1* from 12-day-old plants grown in 8L:16D treated with or without 90 mM sorbitol or sucrose at ZT12. Samples were harvested at ZT16. In both a and b, *UBQ10* was used as the internal control. Different letters indicate significant differences as determined by one-way ANOVA followed by Dunnett’s T3 multiple comparison test; p≤0.05. Error bars indicate SD (n = 3). **c**, qRT-PCR of *MIPS1* and *FT* from 12-day-old wild-type (Col) and *pgm-1* mutant grown in 16L:8D and 8L:16D conditions. *UBQ10* was used as internal control. Error bar indicates SD. *, *p* ≤ 0.05; **, *p ≤* 0.01; ***, *p ≤* 0.001 (Welch’s t-test). n = 3. **d**, Representative images and shoot fresh weight of wild-type (Col), *mips1*, *pgm-1*, and *pgm-1 mips1* mutants grown in16L:8D (28 days) and 8L:16D (8 weeks) with light intensity of 200 μmol/m^2^ s. **e**, Representative images and shoot fresh weight of Col, *mips1*, and *MIPS1* over-expression lines (MIPS1 OX # 2-1 and # 4-2) grown in 16L:8D (28 days) and 8L:16D (8 weeks) with light intensity of 200 μmol/m^2^ s. In both d and e, different letters indicate significant differences as determined by one-way ANOVA followed by Dunnett’s T3 multiple comparison test; p ≤ 0.05. Error bars indicate SD (n = 5). Scale bar = 1 cm. Raw data and calculations for physiology experiments are presented in Supplementary Table 2.

Photoperiod-controlled sucrose levels rely on the regulated sequestration of sucrose as the storage sugar, starch ^10,20–27^. Furthermore, starch production is necessary for proper daylength measurement in the MDLM system. The starchless mutant, *phosphoglucomutase (pgm-1*), is unable to produce starch or regulate sucrose levels in a photoperiodic manner, causing a malfunction of MDLM ^8,28^. We measured expression of *MIPS1* in the *pgm-1* mutant in 16L:8D and 8L:16D (Fig. 3c). In 16L:8D *MIPS1* expression is altered with a greater amplitude at the ZT4 peak of expression and a less pronounced peak at ZT16 in *pgm-1* as compared to the wide type. Strikingly, in 8L:16D *MIPS1* expression has a much larger amplitude of expression during the light period in *pgm-1* showing inability to maintain low levels of *MIPS1* in short days. We tested the genetic relationship between *PGM* and *MIPS1* by crossing the *mips1-2* mutant with the *pgm-1* mutant and assaying fresh rosette weight in 16L:8D and 8L:16D (Fig. 3d and Supplementary Fig. 3a). We found that the *pgm-1 mips1-2* double mutant shows greater defects in rosette fresh weight in 8L:16D than the *pgm-1* mutant alone. This correlates with the observation that *MIPS1* expression is improperly induced in 8L:16D in the *pgm-1* mutant.

It was previously shown that myo-inositol is not sufficient to promote growth on its own^11^. We confirmed this by generating two transgenic lines that overexpress *MIPS1* (Supplementary Fig. 3b), growing the plants in 8L_200_:16D, and measuring their fresh weight (Fig. 3e and Supplementary Fig. 3c). As expected, the *MIPS1-OX* lines did not have greater fresh weight on average than the wild-type plants. In support of this, myo-inositol supplementation in 8L_100_:16D also did not promote growth (Supplementary Fig. 3d). Together with the results in the *pgm-1* mutant, this suggests that *MIPS1* and myo-inositol production are required for rapid long day growth but not sufficient for rapid growth in short days. This indicates that there are other MDLM or photoperiod-controlled processes that must be co-activated with *MIPS1* to promote growth, or alternately that there is a growth-restrictive mechanism that must be suppressed for *MIPS1* to promote growth.

Our results show that the CO/FT mechanism has little or no effect on *MIPS1* expression or photoperiodic growth under the conditions tested here. Interestingly, the *pgm-1* mutant shows defects in flowering time, but published results suggest that *FT* is not misregulated in the *pgm-1* mutant in 12L:12D ^29^. To further test this, we measured *FT* expression in the *pgm-1* mutant in 8L:16D and 16L:8D (Fig. 3c). *FT* expression in the *pgm-1* mutant is the same as wild type in 8L:16D and 16L:8D, suggesting that the MDLM and CO photoperiod measuring systems control distinct target genes to regulate growth and flowering.

### MIPS1 activity and plant growth are determined by the length of the photosynthetic period

The CO/FT and MDLM/MIPS1 photoperiod regulons function in parallel with respect to the molecular components that comprise each mechanism. Next, we wanted to use the *mips1* mutant growth defects to test if the two systems measure the same photoperiod. The CO/FT system uses blue and red light photoreceptors to measure the duration of the day, even at very low light levels, a near “absolute” daylength ^30^. Our results indicate that the photosynthetic complex is required for MDLM to control induction of *MIPS1* expression in long days, indicating that Arabidopsis may also be able to measure the photosynthetic period in parallel to the absolute photoperiod. To test this, we started by reducing light intensity to 40 μmol/m^2^ s, which is near or below the light compensation point (light intensity where the rate of photosynthesis matches the rate of cellular respiration) for Arabidopsis ^27^. At the compensation point, red and blue light photoreceptors remain active, but photosynthesis is unable to produce more sugar than is used by respiration, leaving little extra usable sugar for processes such as growth. In 16L_40_:8D, the *mips1-2* mutant shows no defects in growth, flowering, or fitness when compared to wild-type plants indicating that *MIPS1* is not required for growth in long days when light is below the compensation point (Fig. 4a, b and Supplementary Fig. 4a-c). Conversely, wild-type and *mips1-2* mutant plants flower at approximately 35 days, which is greatly accelerated compared to flowering time in 8L_100_:16D (~82 days)) or 8L_200_:16D (~65 days) (Supplementary Figs. 1a, 4b). Because 16L_40_:8D has a lower daily light integral than these two conditions, this suggests that photoperiodic flowering, which is under the control of photoreceptors that are active at 40 μmol/m^2^ s, is functioning normally.

**Fig. 4:**
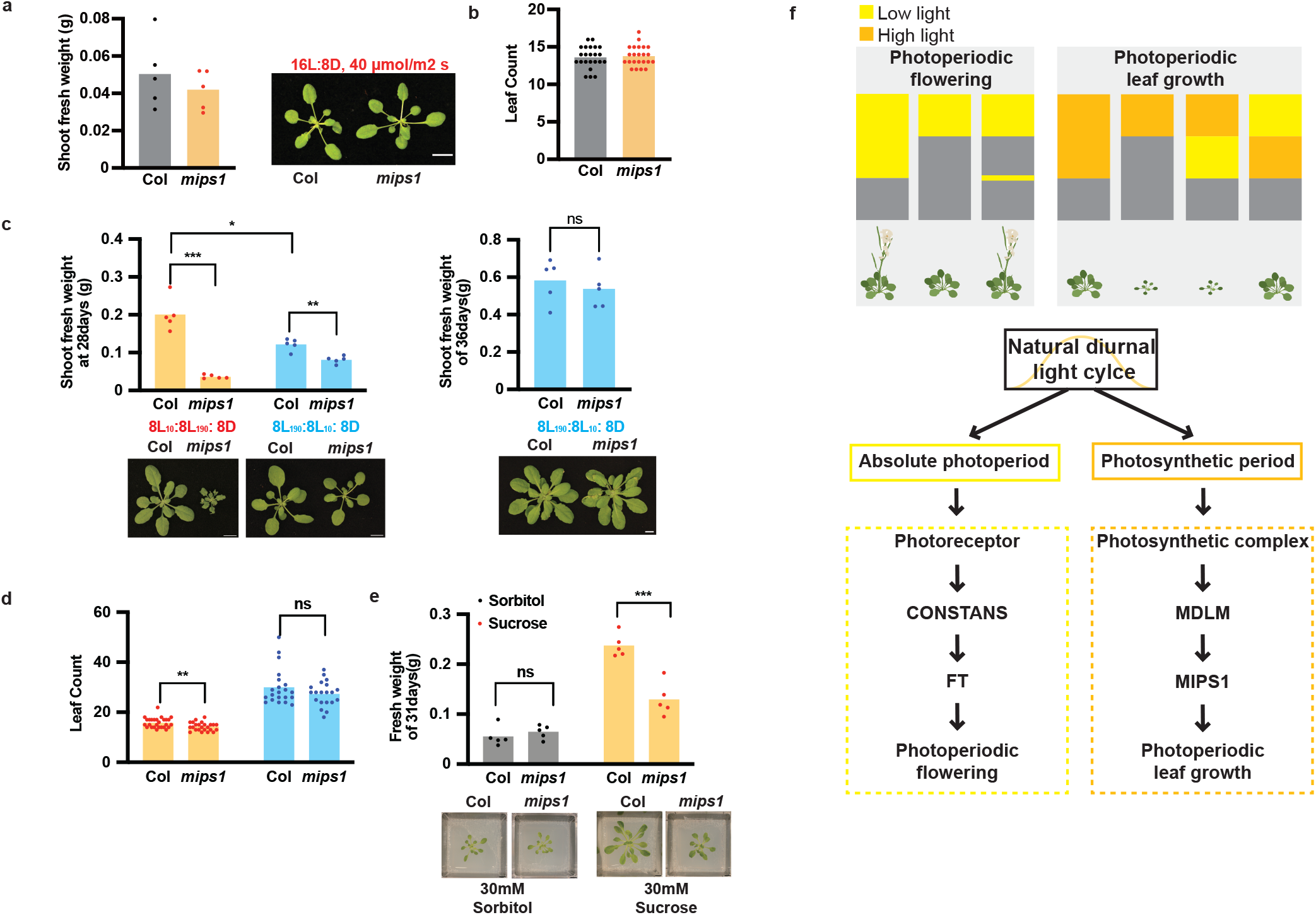
Photosynthetic period controls *MIPS1*-dependent growth. **a** and **b**, Shoot fresh weight (a) with representative image, and leaf number at 1 cm bolting (b) of 28-day-old wild-type (Col) and *mips1* mutant grown in 16L:8D with light intensity of 40 μmol/m^2^ s. No significant difference was detected by Welch’s t-test. Scale bar = 1 cm. n = 5 in a and 23 in b. **c** and **d**, Representative images, shoot fresh weight (c) and leaf number at 1 cm bolting (d) of Col and *mips1* grown in 8L_190_:8L_10_:8D and 8L_10_:8L_190_:8D conditions. n = 5 in c and n = 20 – 27 in d. **e**, Representative images and fresh weight of Col and *mips1* grown in 8L:16D with light intensity of 200 μmol/m^2^ s treated with 30 mM sorbitol or sucrose. Plants were geminated on 1/2 MS plates and grown in 12L:12D for 7 days then transferred to the indicated media for 24 days. n = 5. In c, d and e, Error bar indicates SD. **, *p ≤* 0.01; ***, *p ≤* 0.001 (Welch’s t-test). **f**, Model for two distinct photoperiod measurement mechanisms controlling photoperiodic flowering and leaf growth. Within a natural diurnal light cycle are multiple photoperiods that can be distinguished by plants. The classic model of photoperiod flowering is shown on the left and the model for photoperiodic growth on the right. 24 hour periods are shown with low light in yellow, high light (above the compensation point) in orange, and dark in gray. For photoperiodic flowering, low light daylengths are measured to control flowering, and the light sensitive portion of the day can be determined by a night break experiment. Light or dark are sensed by photoreceptors which control CONSTANS stability which in turn activates *FT* expression to trigger flowering. For photoperiodic growth, high light, or photosynthetic period, is measured to trigger growth. “Fast break” experiments show the high light sensitive portions of the day. Light above the compensation point is sensed by the photosynthetic complex and measured by the MDLM system to control *MIPS1* expression which is required to support rapid leaf growth in long days.

Classic night break experiments show that there are transient periods of light sensitivity, usually in the middle 8 hours (the time of day that has seasonal variation between light and dark) of a 24-hour period, that allow for photoperiod measurement. Here, we found that photoperiodic plant growth is promoted, and the *mips1* phenotype is apparent, in 16L_100_:8D and 16L_200_:8D, but growth is restricted and the *mips1* mutant phenotype is absent in 8L_100_:16D, 8L_200_:16D, and 16L_40_:8D. These results suggest that the middle 8 hours of the day, ZT8-ZT16, is more sensitive to light above the compensation point than the first 8 hours of the day, ZT0-ZT8, with respect to growth. We tested this directly by growing the wild type and *mips1* mutant in two long day lighting regimes, one with high light (190 μmol/m^2^ s) from ZT0-ZT8 and low light (10 μmol/m^2^ s) from ZT8-ZT16 (8L_190_:8L_10_:8D) and one with low light (10 μmol/m2 s) from ZT0-ZT8 and high light (190 μmol/m^2^ s) from ZT8-ZT16 (8L_10_:8L_190_:8D) (Fig. 4c,d and Supplementary Fig. 4d). In these experiments the daily light integral and day length are held constant, but the time of day when the compensation point is crossed and photosynthates are available for growth is altered. Rather than a “night break” experiment, this may be better termed a “fast break” experiment because we are interrupting a low light and low photosynthate long day with light above the compensation point to produce photosynthates during a specific period. The *mips1* mutant plants grown in 8L_190_:8L_10_:8D showed no altered phenotype compared to the wild-type plants when examined at 36 days (time of flowering) and show no lesions but had generated slightly less fresh weight at the 28 day timepoint. Strikingly, the *mips1* plants grown in 8L_10_:8L_190_:8D showed severe growth and cell death phenotypes at 28 days, similar to plants grown in 16L_200_:8D (Figs. 1d and 4c), and we were not able to collect fresh weight data at 36 days because they had flowered at 28 days. Perhaps most importantly, the wild-type plants grew nearly 70% larger in the 8L_10_:8L_190_:8D, suggesting that light above the compensation during ZT8-ZT16, has a greater effect on growth than light during ZT0-ZT8. This indicates that the light sensitive time of day for growth occurs between ZT8 and ZT16 when *MIPS1* is induced in long day photoperiods.

Conversely, flowering time in 8L_190_:8L_10_:8D is earlier than the 8L_200_:8D condition suggesting the flowering pathway is still measuring the absolute daylength. Interestingly, the 8L_10_:8L_190_:8D growth condition induced flowering earlier than the in 8L_190_:8L_10_:8D (Fig. 4d and Supplementary Fig. 4d), suggesting that high light from ZT8-ZT16 can accelerate long day flowering as well. Together these results confirm the idea that photoperiodic flowering is controlled by low light, a near “absolute daylength”, but growth is controlled by the photosynthetic period and that these two photoperiods are dissociable (Fig. 4f).

The “fast break” experiments indicate that the plant is likely responding differentially to the presence or absence of photosynthates at ZT8-ZT16. To test this, we performed sucrose supplementation experiments in a short-day growth condition which should aberrantly promote long day-like growth and thus require the sucrose-induced expression of *MIPS1* (Fig. 3b). We grew wild-type and *mips1* mutant plants in 8L_200_:16D, where growth is normally restricted and *MIPS1* is not required, in the presence of sucrose or the control non-hydrolyzable sugar, sorbitol (Fig. 4e). Sucrose promoted rapid growth in the wild-type plants, and this growth was diminished in the *mips1* mutant. This shows that plants are detecting the photosynthetic period by timing sensitivity to sugar production, resulting in a metabolic daylength, and that promotion of growth by photosynthetic sugars at this time requires *MIPS1*.

## Discussion

This work provides a new view of seasonal timekeeping in plants by showing that photoperiodic growth is controlled by a wholly separate molecular mechanism from photoperiodic flowering, despite often occurring in the same season (Fig. 4f). We accomplished this by defining one, of likely multiple, long day photoperiodic growth pathways, from photoperiod detection to gene expression. This photoperiodic growth pathway relies on the MDLM system to convert photosynthetic period into a duration of photosynthate production that plants can detect to control transcription of genes required for growth, such as *MIPS1*.

Classic flowering mutants, such as *ft* or *co*, with easily observable defects in flowering time precipitated the discovery of the photoperiodic flowering mechanism. In a similar way, our studies relied on the easily observed growth defects of the *mips1* mutant to uncover a mechanism for photoperiodic growth. The *mips1* mutant is small and forms lesions, exclusively in long days allowing us to easily detect the *mips1* phenotype as an indicator for long day photoperiod. In addition, *MIPS1* is expressed with an easily distinguishable photoperiod-specific expression pattern, similar to *FT*.

The usefulness of the *mips1* mutant is shown in the “fast break” (Fig. 4c, d) and sucrose supplementation experiments (Fig. 4e) which clearly demonstrate that Arabidopsis is measuring photosynthetic period and is integrating this information as the duration of time that photosynthetic sugars are being produced, a metabolic daylength. This demonstrated that growth is truly a photoperiodic process but that the type of photoperiod that is detected for growth, photosynthetic period, is different than that detected for flowering, which is a near absolute photoperiod perceived by low light photoreceptors^30^. It is remarkable that photosynthetic period is measured in parallel to absolute photoperiod and suggests different types of day lengths enact distinct gene expression networks. This is the simplest explanation for the disconnect between photoperiodic growth and flowering in short day plants that was recognized a century ago^6^. This also shows that that natural diurnal light cycles do not constitute a single photoperiod, but rather are composed of many different photoperiod types that can be differentially detected to independently coordinate many different seasonal developmental processes.

## MATERIALS AND METHODS

### Arabidopsis materials

The *Arabidopsis* seeds of Col, *mips1-2* (SALK_023626), *pgm-1* (CS210), *co-9* (CS870084), *ft-10* (CS9869) were obtained from ABRC. The *mips1-2* mutant was also crossed to *co-9* and *pgm-1* mutant plants and the *co-9 mips1-2* and *pgm-1 mips1-2* double mutants were identified by genotyping. The *pgm-1* allele was genotyped as described before ^31^. The primers used for genotyping are listed in Supplementary Table 1.

### Arabidopsis growth conditions

*Arabidopsis* thaliana seeds from Col and mutants were surface sterilized for 20 min in 70% ethanol with 0.1% Triton X-100 then sown on freshly poured 1/2 MS plates, pH 5.7, (Cassion Laboratories, cat. # MSP01) and 0.8% bacteriological agar (AmericanBio cat. # AB01185) without sucrose. The seeds were stratified in the dark for two days at 4 °C then transferred into 22 °C, 12L:12D illuminated by white fluorescent lamps at 150 μmol/m^2^ s for seven days. The seven-day-old seedlings were then transferred to different photoperiod for given experiments as indicated. For sucrose treatment, seedlings transferred to culture vessels (vented, PhytoTech) contained 1/2 MS with 30mM Sorbitol or Sucrose. For soil grown plants, after two days stratification, seeds were germinated and grown in Premier Horticulture PRO-MIX BX at 22°C in 16L:8D or 8L:16D with light intensity of 10/40/100/190/200 μmol/m^2^ s as indicated. For Myo-inositol (Sigma-Aldrich) treatment, the plant was sprayed with 50mg/mL myo-inositol solution (dissolute in water) or water every 2 days^11^.

### Plasmid construction

For the *MIPS1* overexpression plasmids, the *MIPS1* coding sequence was obtained by PCR using Col cDNA as the template, inserted into pENTR/D-TOPO and then transferred into PB7-HFN destination vectors using LR recombination ^32^.

### qRT-PCR

For qRT-PCR experiments, RNA extraction was performed with two different methods. For Fig. 1b and 1c, total RNA was extracted from Arabidopsis seedlings grown in indicated conditions using TRIzolTM reagent (ThermoFisher, cat. # 15596026); for the remaining Figs. 2a, 3a-c and supplementary fig. 3b extraction was performed with RNeasy Plant Mini Kit (QIAGEN cat. # 74904). In both methods, the resulted RNA was subsequently treated with DNase (QIAGEN, cat. # 79254). The subsequent reverse-transcription and conditions for qRT-PCR reactions were described previously with minor modifications ^33^. Briefly, four hundred nanograms of total RNA were used for reverse-transcription using iScript Reverse Transcription Supermix for RT-qPCR (Bio-Rad, cat. # 1708841). iTaq Universal SYBR Green Supermix was used for qRT-PCR reaction (Bio-Rad, cat. # 1725121). *IPP2* (AT3G02780) or *UBQ10* (AT4G05320) was used as an internal control as indicated. The relative expression represents means of 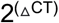 from three biological replicates, in which ΔCT = (CT of Gene of Interest – CT of internal control). The primers used are listed in Supplementary Table 1.

## ACKNOWLEDGEMENTS

We would like to thank Christopher Adamchek for technical support. We would also like to thank Sandra Pariseau and Jenny Pengsavath for administrative support. Additionally, we would like to thank Chris Bolick, Nathan Guzzo, and the staff at Marsh Botanical Gardens for their support in maintaining plant growth spaces. We would also like to thank Dr. Adam Saffer, Lilijana Oliver, Harper Lowrey, Damon Clark, and Shirin Bahmanyar for insightful discussions and critical reading of the manuscript. This work was supported by the National Institutes of Health (R35 GM128670) to J.M.G. W.L. and Q.W. were supported by the Forest BH and Elizabeth DW Brown Fund Fellowship. DT was supported by the National Institutes of Health (T32GM007223-44).

## AUTHOR CONTRIBUTIONS

Q.W., T.L., W.L., and J.M.G. designed the experiments. Q.W., T.L., W.L., and D.T. performed the experiments and experimental analyses. Q.W. and J.M.G. wrote the article.

## Figure Legends

**Supplementary Fig. 1:**
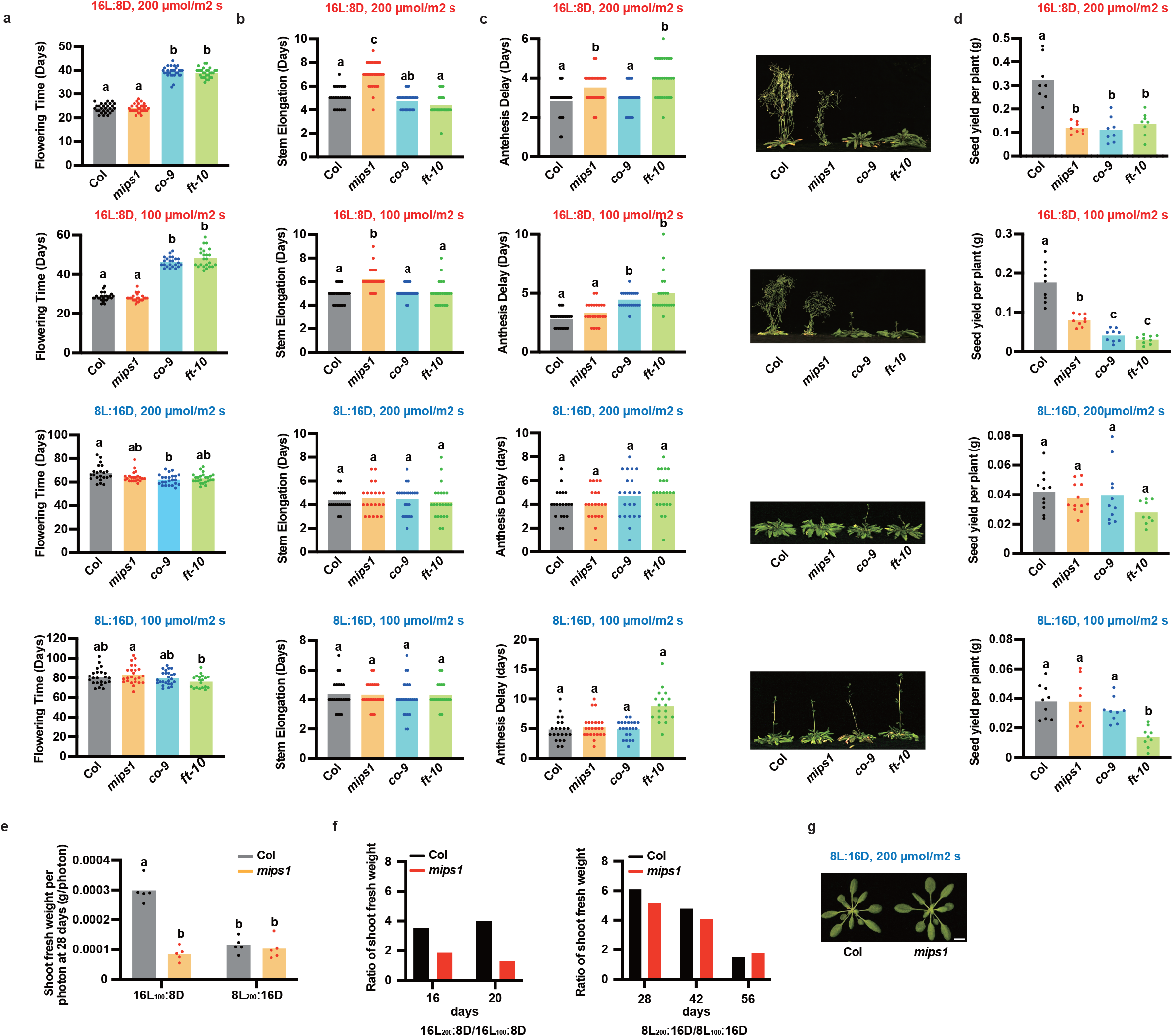
*MIPS1* is involved in light intensity-dependent and photoperiod-dependent leaf growth. **a-d**, Flowering time (a), stem elongation time (b), anthesis delay (c) with representative images, and yield (d) of Col, *mips1, co-9* and *ft-10* plants grown in 16L:8D or 8L:16D conditions with light intensity of 200 μmol/m^2^ s or 100 μmol/m^2^ s. n = 8-30. **e**, Shoot fresh weight per day per available photon of 28-day-old Col and *mips1* plants grown in 16L:8D or 8L:16D conditions with light intensity of 100 μmol/m^2^ s or 200 μmol/m^2^ s as indicated. n = 5. **f**, Ratio of shoot fresh weight of Col and *mips1* plants grown in16L_200_:8D comparing to16L_100_:8D and 8L_200_:16D compared to 8L_100_:16D at indicated age. **g**, Image of 28-day-old Col and *mips1* plants grown in 8L:16D condition with light intensity of 200 μmol/m^2^ s. In a, b, c, d and e, different letters indicate significant differences as determined by one-way ANOVA followed by Dunnett’s T3 multiple comparison test; p≤0.05. Raw data and calculations for physiology experiments are presented in Supplementary Table 2.

**Supplementary Fig. 2:**
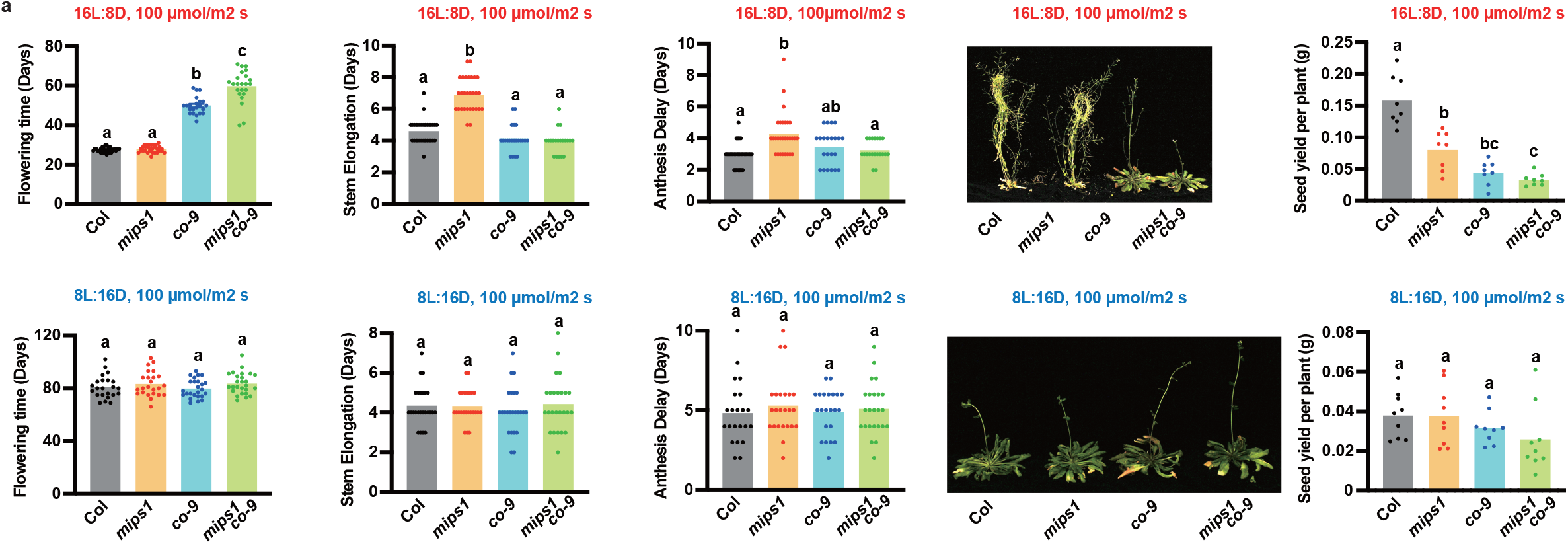
*MIPS1* function is distinct from CO/FT. **a,** Flowering time, stem elongation time, anthesis delay with representative images, and yield of Col, *mips1, co-9* and *mips1 co-9* plants grown in 16L:8D or 8L:16D conditions with light intensity of 100 μmol/m^2^ s. Different letters indicate significant differences as determined by one-way ANOVA followed by Dunnett’s T3 multiple comparison test; p≤0.05. Error bars indicate SD (n = 8-30). The data were re-plotted from Fig. S1a. S1b, S1c and S1d, respectively. Raw data and calculations for physiology experiments are presented in Supplementary Table 2.

**Supplementary Fig. 3:**
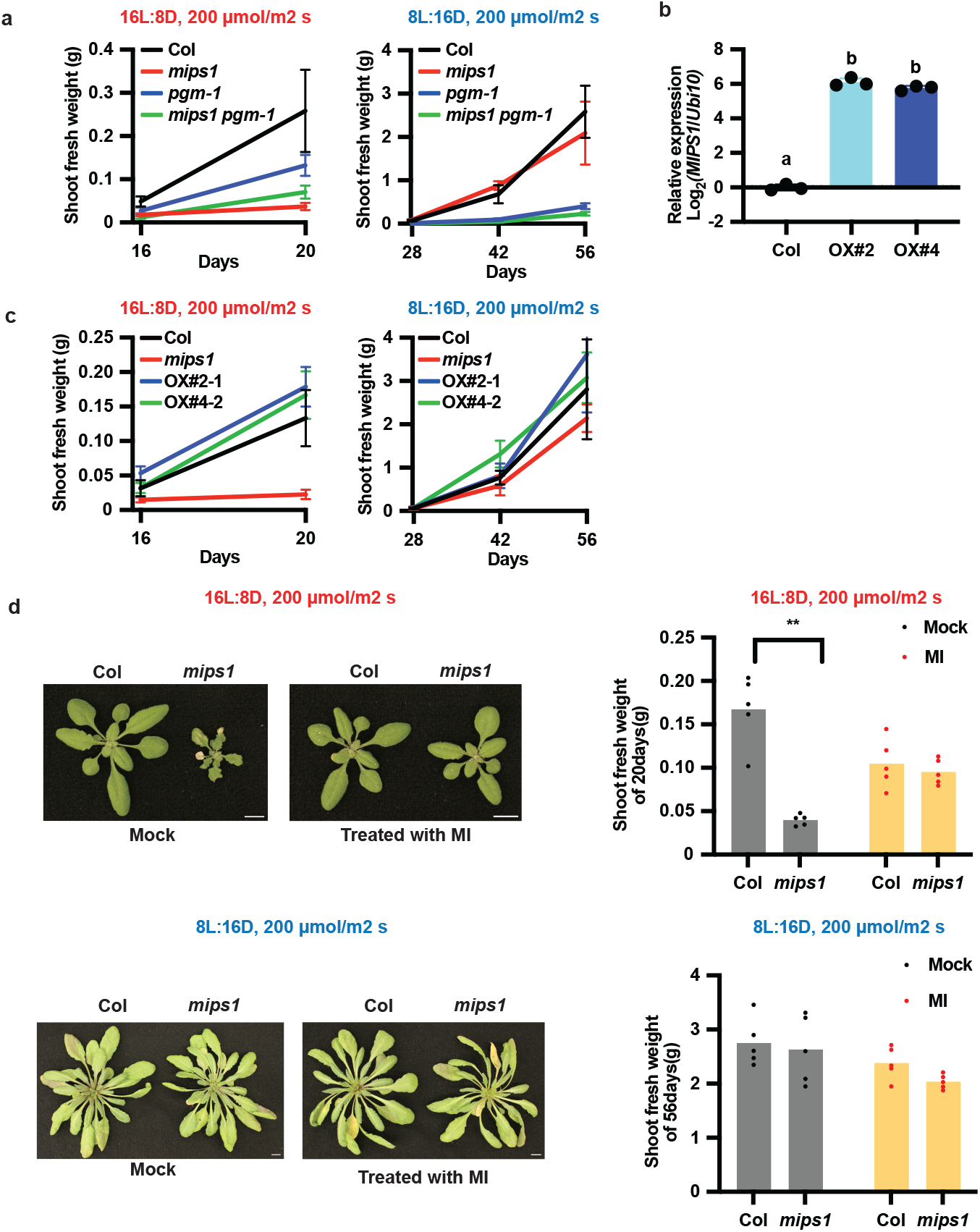
Myo-inositol is required for, but not sufficient to, induce rapid growth. **a**, Time course of fresh weight of Col, *mips1, pgm-1* and *pgm-1 mips1* grown in16L:8D or 8L:16D condition with light intensity of 200 μmol/m^2^ s (n = 5). **b**, qRT-PCR of *MIPS1* from 12-day-old Col, MIPS1OX # 2-1 and # 4-2 plants grown in 8L:16D. Samples were harvested at ZT12. *UBQ10* was used as internal control. Different letters indicate significant differences as determined by one-way ANOVA followed by Dunnett’s T3 multiple comparison test; p≤0.05. Error bars indicate SD (n = 3). **c**, Time course of fresh weight of Col, *mips1*, MIPS1OX # 2-1 and # 4-2 plants grown in16L:8D or 8L:16D condition with light intensity of 200 μmol/m^2^ s (n = 5). **d**, Representative images and shoot fresh weight of Col and *mips1* grown in 16L:8D (20 days) and 8L:16D (8 weeks) with light intensity of 200 μmol/m^2^ s sprayed with 50 mg/mL myo-inositol solution or water every 2 days. Error bar indicates SD. **, *p* ≤ 0.01 (Welch’s t-test). n = 3.

**Supplementary Fig. 4:**
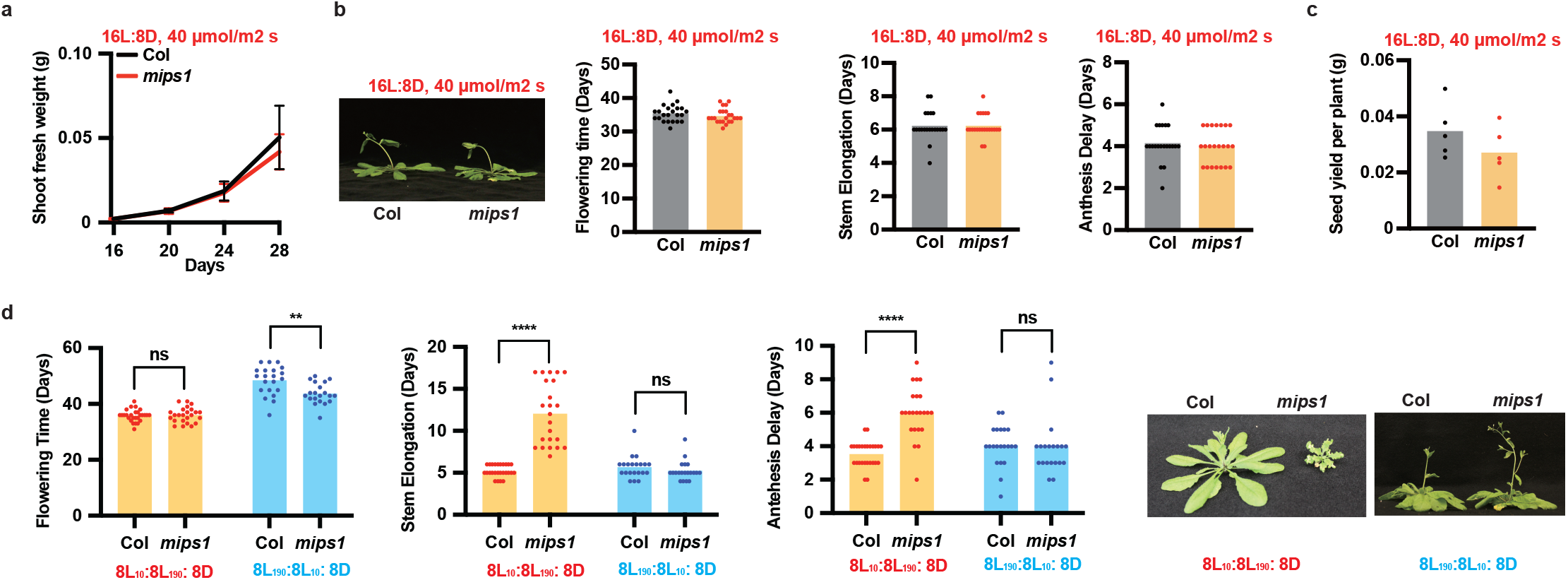
Photosynthetic period controls *MIPS1*-dependent growth. **a-c**, Time course of fresh weight (a), flowering time, stem elongation time, anthesis delay (b) with representative images, and seed yield (c) of Col and *mips1* plants grown in16L:8D with light intensity of 40 μmol/m^2^ s. n = 5 in a, n = 21-23 in b, and n = 5 in c. Error bar indicates SD. No significant difference was detected by Welch’s t-test. **d**, Flowering time, stem elongation time and anthesis delay of Col and *mips1* plants grown in 8L_190_:8L_10_:8D and 8L_10_:8L_190_:8D conditions. Error bars indicate SD. **, *p* ≤ 0.01; ****, *p* ≤ 0.0001 (Welch’s t-test). n = 20-27. Raw data and calculations for physiology experiments are presented in Supplementary Table 2.

**Supplementary Table 1:**
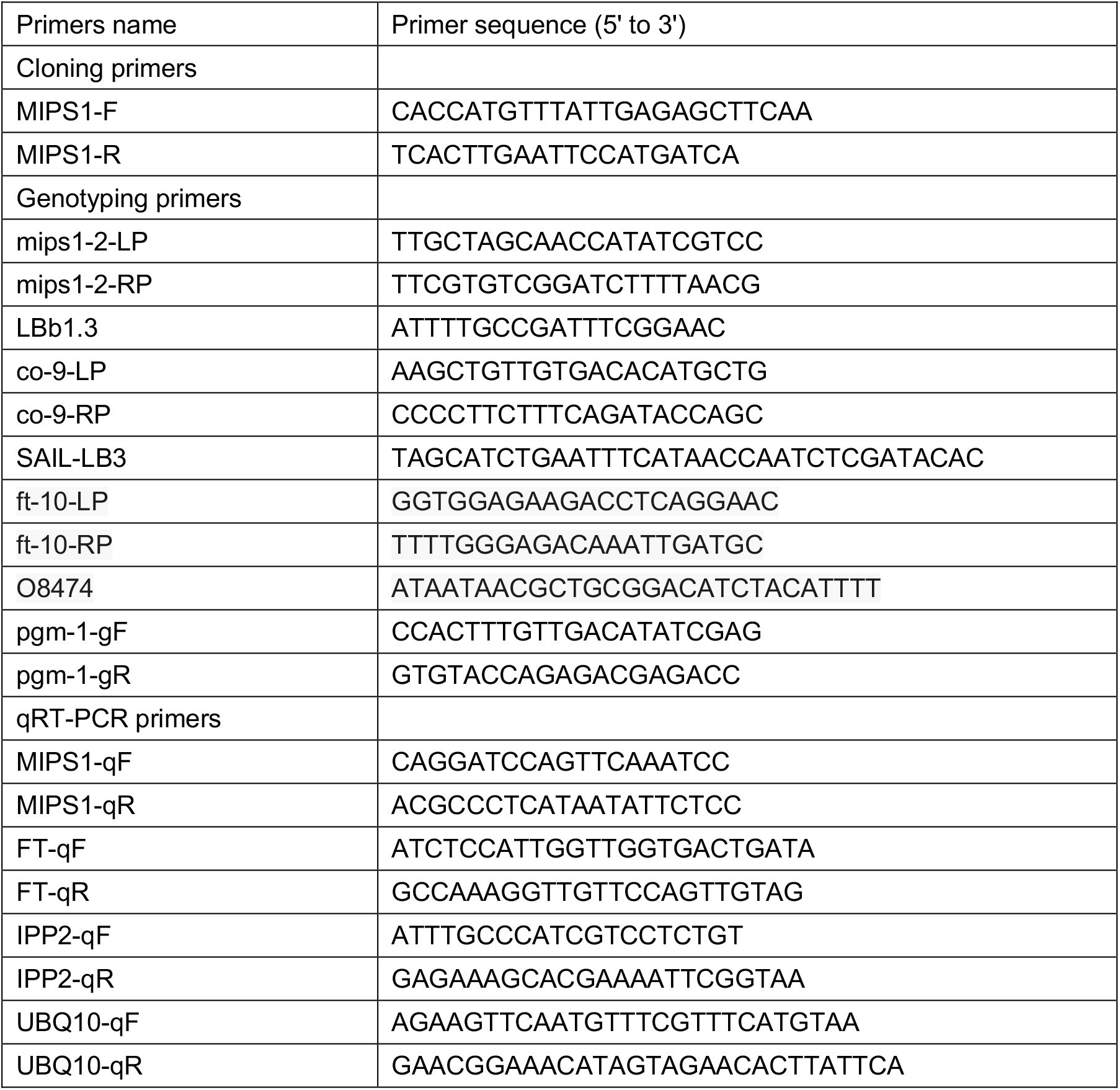
Primers used in this study

